# UniFacePoint-FM: A Foundation Model for Generalizable 3D Facial Representation Learning and Multi-Attribute Prediction

**DOI:** 10.64898/2026.02.06.703926

**Authors:** Dan Li, Chih-Hsiung Fu, Kun Tang

## Abstract

The human face is a rich medium for biometric, behavioral, and clinical information. However, 2D facial images based technologies lack critical geometric details and are susceptible to pose and illumination interference, while 3D facial deep learning frameworks are hindered by complex annotation, preprocessing, and task-specific designs with poor cross-domain generalization. To address these challenges, we propose UniFacePoint-FM, a 3D facial foundation model built on a self-supervised Point-MAE framework, tailored for high-fidelity point cloud representation learning. The model was pretrained on a self-constructed dataset of high-resolution 3D facial scans, followed by supervised fine-tuning and comprehensive evaluation across three independent datasets for diverse downstream tasks. Experimental results demonstrate that UniFacePoint-FM is both pretraining-efficient and highly generalizable: it achieves state-of-the-art performance on gender classification, age regression, and BMI prediction, and matches the accuracy of the ResMLP model (while outperforming other baselines) in facial expression recognition. Notably, by learning high-quality, fine-grained representations directly from raw point clouds, UniFacePoint-FM delivers robust generalization and transferability across tasks, datasets, and even different face scanning platforms. Overall, our work establishes an effective foundation model paradigm for 3D facial analysis, with promising implications for biometric security, health monitoring, and advanced human-computer interaction systems.

## 1. Introduction

The human face serves as a rich carrier of diverse information, encompassing biometric traits (e.g., identity, gender, age, ethnicity, BMI), behavioral responses (including facial expressions and the underlying emotional states), as well as health and clinical cues[1, 2, 3, 4]—with numerous genetic syndromes linked to craniofacial abnormalities[5, 6]. This informational value underscores the pivotal role of facial analysis in computer vision. Meanwhile, face image/model generation techniques have become increasingly powerful and versatile, facilitating the creation of virtual reality characters[7, 8]. Computer vision in human face has boasted extensive applications in security (e.g., surveillance), healthcare (e.g., mental health monitoring), entertainment, and human-computer interaction. Traditionally, facial analysis—spanning classification, recognition, and generation—has relied heavily on 2D imagery, where pixels are processed within a regular grid format. Deep learning architectures, most notably Convolutional Neural Networks (CNNs), Variational Autoencoder(VAE) and Transformers, have been extensively optimized for these 2D applications, achieving unprecedented accuracy and representational capacity. However, as highlighted by Fei-Fei Li, the physical world is inherently three-dimensional, and “spatial intelligence” is required for AI to perceive and interact with the world in its native form [From Words to Worlds: Spatial Intelligence is AI’s Next Frontier]. 2D facial images are merely flattened projections of 3D structures, often confounded by extrinsic variables such as pose, illumination, and cosmetic textures. This dimensionality loss discards vital geometric information, spatial relationships, and dynamic curvature changes, which can lead to suboptimal performance in nuanced tasks like micro-expression classification, emotion recognition, and clinical disease detection, etc. The shift toward deep learning approaches that directly process 3D facial analysis represents a burgeoning frontier. Despite its promise, 3D facial analysis faces two primary bottlenecks: Data Scarcity and Annotation Complexity. Acquiring largescale, high-quality 3D datasets is logistically intensive and costly. Accurate biometric or demographic labeling often requires specialized equipment and controlled environments. However, this gap is narrowing due to the rapid evolution of 3D imaging technologies, including structured light, Time-of-Flight (ToF) sensors, and high-resolution 3D LiDAR and Millimeter-Wave (mmWave) radar[9, 10, 11, 12]. Secondly, in contrast to the regular pixel grids of 2D images, 3D face data typically exists as irregular, unordered, and non-uniformly sampled point clouds[13]. Consequently, traditional pipelines often require elaborate preprocessing—such as mesh generation and landmarking and dense alignment—before feature learning can begin. Although numerous strategies have been proposed to address these topological irregularities, they often incur significant computational costs that hinder real-time application. Due to these constraints, deep learning research in the 3D facial domain remains relatively limited; existing approaches are often developed in a task-specific manner, exhibiting considerable room for improvement in predictive accuracy while struggling to generalize across task, datasets, and domains. Li *et al*.[14] attained a 95.4% gender classification accuracy using a 3D facial deep learning model. Chen *et al*.[15] reported an age prediction mean absolute error of approximately 6 years. Point cloud preprocessing also frequently degrades performance in cross-domain transfer. Therefore, a robust and efficient deep learning framework may be achieved via direct encoding of raw facial point clouds, eliminating extra alignment and landmarking steps.

In recent years, transformer-based point cloud modeling has been developed to directly tackle 3D object recognition from unaligned raw point clouds [16, 17, 13]. Following the foundation model paradigm, these models enable effective scaling via self-supervised pretraining and can be fine-tuned to adapt to a wide range of tasks. Among these, Masked Autoencoders (MAE) enables learning robust geometric features by reconstructing masked input subsets[18]. Extending MAE to facial recognition is intuitive yet challenging. Unlike general 3D objects with distinct inter-class geometric variations, human faces feature subtle, fine-grained shape differences and much higher point densities[19, 20], making 3D facial analysis a highly specialized task.

In this study, we introduce UniFacePoint-FM, a large-scale 3D facial foundation model built upon the Masked Autoencoder (MAE) paradigm and optimized for high-fidelity point cloud representation learning. Leveraging the self-supervised Point-MAE framework[13], this model is pretrained on thousands of high-resolution 3D facial scans. Post pretraining, we perform fine-tuning on multiple independent datasets across diverse tasks, including gender classification, age regression, BMI prediction, and facial expression recognition. Experiments show UniFacePoint-FM is pretraining-efficient and highly generalizable, achieving state-of-the-art performance in nearly all facial attribute recognition tasks.

In summary, this work provides the following primary contributions:

- We introduce UniFacePoint-FM, a foundation model specifically tailored for generalizable 3D facial representation learning based on Point clouds.
- We perform systematic multi-dataset and multi-task evaluations, demonstrating strong cross-task and cross-domain generalization.
- We show that UniFacePoint-FM supports accurate prediction of multiple facial attributes and achieves competitive or superior performance across benchmarks.
- We establish an effective and scalable paradigm for 3D facial foundation models, offering broad potential for biometric security, health monitoring, and human–computer interaction.

## 2. Related Work

### 2.1. Self-Supervised Pretraining in NLP and 2D Vision

Self-supervised learning has reshaped representation learning in NLP and 2D vision by enabling models to learn semantic and structural information directly from unlabeled data. In NLP, masked language modeling (MLM) frameworks such as BERT, RoBERTa, and ALBERT have demonstrated that reconstructing masked tokens from context yields highly transferable linguistic representations[21, 22, 23]. Large-scale generative self-supervised pretraining, exemplified by GPT-3, further extended this paradigm by enabling powerful zero-shot and few-shot generalization across tasks[24]. In 2D computer vision, ViT bridged the architectural gap between vision and NLP by treating images as patch sequences[25]. This unified framework laid the foundation for Masked Image Modeling (MIM) methods like MAE[26] and SimMIM[27], enabling efficient representation learning via BERT-style masked reconstruction, and related models introduced masked patch reconstruction as a simple yet effective pretext task, establishing strong baselines for downstream tasks such as classification, detection, and segmentation.

### 2.2. Self-Supervised Representation Learning for 3D Point Clouds

Recent research has concentrated on masked modeling for point clouds. Point-BERT introduced a discrete tokenization strategy based on Farthest Point Sampling (FPS) and reconstructs masked tokens within a BERT-style framework[28]. Point-MAE further advances this paradigm by proposing a fully geometric masked autoencoder featuring an asymmetric design, which utilizes a standard Transformer encoder and a lightweight decoder to directly reconstruct masked point patches[13]. This enables Point-MAE to achieve superior performance on 3D object classification and segmentation benchmarks. Subsequent models, such as Point-M2AE, incorporate refined multiscale masking strategies and pyramidal architectural improvements, further enhancing the quality of learned 3D representations and achieving consistent gains across downstream benchmarks[29].

### 2.3. 3D Human Facial Attribute Prediction

Research on 3D facial analysis encompasses face recognition, demographic attribute estimation, expression recognition, and various medical or biometric assessments. Traditional methodologies rely on hand-crafted geometric descriptors, including curvature measures, depth histograms, and local shape descriptors, typically coupled with classical machine learning algorithms such as Support Vector Machines (SVMs) [30, 31, 32], Partial Least Squares Regression (PLSR) [15], and Random Forest (RF)[33]. To model global shape variation, 3D Morphable Models (3DMMs) have also been widely adopted[34]. Recently, geometric deep learning methods have emerged; for instance, Graph Neural Networks (GNNs) have been utilized for gender classification, Variational Autoencoders (VAEs) have been employed for interpretable attribute analysis, and Convolutional Mesh Autoencoders, such as CoMA, have demonstrated superior performance in facial generation and expression modeling[35, 36, 37]. Furthermore, while Masked Autoencoders (MAEs) have primarily exhibited robust representation learning capabilities in general 3D point cloud understanding, their self-supervised learning paradigm presents new potential for 3D facial analysis.

## 3. Materials and Methods

### 3.1. Dataset

#### 3.1.1. The TaiZhou dataset (TZ)

The TaiZhou dataset was collected in TaiZhou City, China, and comprises 2,977 high-resolution 3D facial scans, among which 2,499 are accompanied by demographic attributes, including gender, age, and body mass index (BMI). As shown in Fig. S1(a), the dataset exhibits a moderately unbalanced gender distribution, with 64.17% females and 35.83% males. The age ranges from 31 to 80 years with an average of 55.84 years, and the BMI ranges from 14.69 kg/m^2^to36.91 kg/m^2^ with a mean of 24.39kg/m^2^.

#### 3.1.2. The ChenZhou dataset (CZ)

The ChenZhou dataset was collected in ChenZhou city, China, comprising 1732 participants who have provided high-resolution 3D facial scans and corresponding demographic data, including sex, age, and BMI. As shown in Fig. S1(b), the majority is female (70.61%), the age range is from 17 to 32 years old, with an average of 19.75 years, and the BMI ranges from 14.68 kg/m^2^ to 33.30 kg/m^2^ with an average of 20.51 kg/m^2^.

#### 3.1.3. The FaceScape dataset (FS)

The FaceScape dataset is a publicly available collection consisting of 847 subjects, each with 20 high-resolution 3D facial scans captured under distinct facial expressions. All 3D scans were acquired using a dense 68-camera array under controlled illumination, followed by multi-view stereo reconstruction. After excluding samples with missing information, a total of 823 subjects were retained for this study. As shown in Fig. S1(c), the gender distribution is nearly balanced, with 50.06% females and 49.94% males. The participants’ age ranges from 18 to 69 years, with an average of 28.5 years. Information on BMI is not available for this dataset.

#### 3.1.4. Ethics statement and consent to participate

All participants provided written informed consent to participate in the project.Sample collection for this study was conducted with the approval of the ethicscommittee of Shanghai Institutes for Biological Sciences and in accordance with thestandards of the Helsinki Declaration.

### 3.2. Data Preprocessing

The 3D images in both the TaiZhou dataset and ChenZhou dataset were acquired using a 3dMDface system[38]. Participants were asked to keep a neutral facial expression during data acquisition. Non-rigid dense registration was performed to align all facial scans to a common template and establish point-wise correspondence using FRAS(Facial Registration Analysis Software) from paper[39]. This process ensured a unified topological structure across all 3D facial data, facilitating subsequent analysis and model training. Crucially, no normalization was performed, ensuring that all scans were maintained in their original scale and orientation.

For the FaceScape dataset, the facial surface was extracted by cropping topology-consistent full-head 3D mesh. Unlike TaiZhou dataset and ChenZhou dataset, where each face contains 32,251 points, the facial scans in this dataset contain only 12,296 points.

### 3.3. UniFacePoint-FM

The UniFacePoint-FM is a model built upon the Point-MAE architecture but is tailored to the structural characteristics of 3D facial point clouds. Fig. 1 illustrates the overall scheme of our approach. The model training consists of two stages: self-supervised learning on unlabeled facial scans and supervised fine-tuning on task-specific data.

**Fig. 1:**
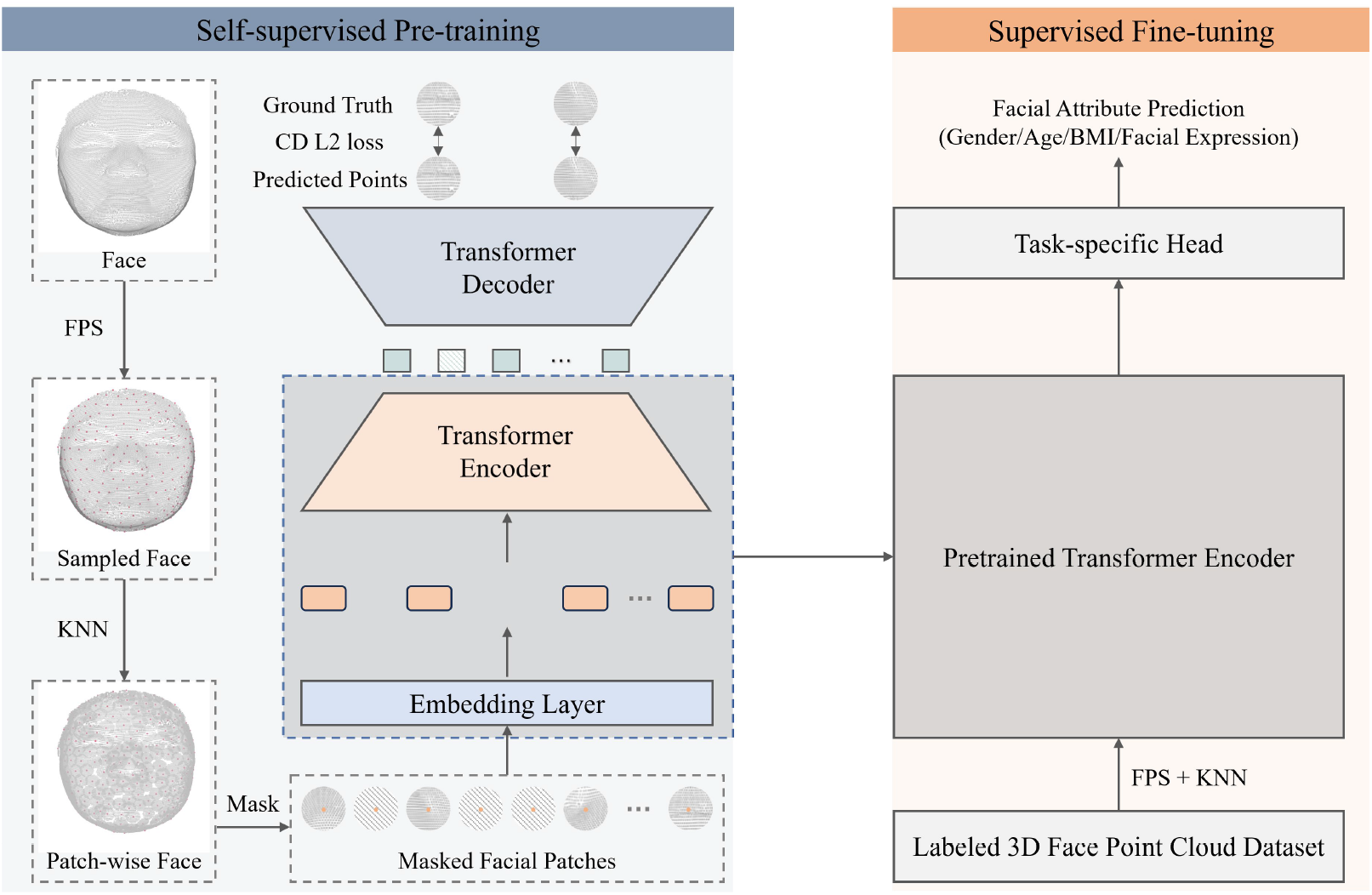
Overview of the proposed self-supervised pre-training and supervised fine-tuning framework for 3D facial representation learning.

#### 3.3.1. Self-supervised Pre-training

We followed the novel and streamlined self-supervised learning strategy introduced in the Point-MAE model.[17]. A 3D face point cloud is first divided into irregular point patches P by applying Farthest Point Sampling (FPS) to select representative centroids, followed by constructing neighborhoods via the K-Nearest Neighbors (KNN) algorithm. Subsequently, 60% of these point patches are randomly masked. The masked point patches*P*_*m*_ are processed as mask tokens *T*_*m*_ by a share-weighted learnable mask token. The visible point patches *P*_*v*_ are embedded and fed into an encoder consisting of standard Transformer blocks to generate encoded tokens *T*_*e*_. The decoder, which is similar to the encoder but contains fewer Transformer blocks, reconstructs masked point patches.

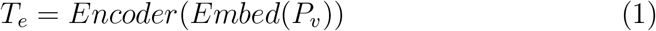

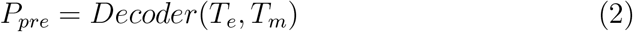

We use *l*_2_ Chamfer Distance as the reconstruction loss L,formulated as:

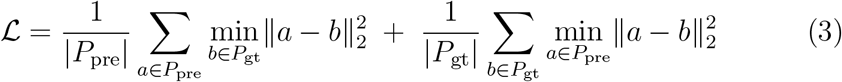

where *a* and *b* indicate the 3D coordinates of points within the predicted point patches *P*_*pre*_ and the ground truth point patches*P*_*gt*_.

It is worth noting that the original Point-MAE is designed for object-level point clouds containing only 1024 points. In contrast, our 3D facial scans have 32,251 points which are significantly denser. The high point density is essential because, unlike generic 3D objects whose overall shapes can be preserved with sparse sampling, subtle geometric variations between different human faces require substantially more points. Excessive down sampling would lead to the loss of crucial individualized facial features. To accommodate the high-density nature of 3D facial data, we increase both the number of groups and the group size in the patch partitioning stage. These adjustments ensure that the resulting patches are sufficiently informative for downstream tasks.

#### 3.3.2. Supervised Finetuning on specific tasks

To evaluate our UniFacePoint-FM model, we attach a task-specific head to the encoder of UniFacePoint-FM and fine-tune the corresponding modules with labeled data for various downstream tasks across different datasets. We conduct experiments on four downstream tasks, including gender classification, age estimation, BMI prediction and expression recognition. The loss function is selected according to the task type. Specifically, Cross Entropy Loss is adopted for gender classification and expression recognition, whereas Mean Absolute Error Loss is used for age and BMI prediction.

## 4. Results

### 4.2. Pretraining-step

A total of 2,977 three-dimensional human face scans were collected in TaiZhou city, Jiangsu Province, China, using 3dMD face system, forming TaiZhou Dataset. All raw 3D images were preprocessed into high-density facial point clouds with uniform topology but kept the original orientation and scale (see Materials and Methods). The dataset was divided into training, validation, and test set with an 8:1:1 ratio. We pre-trained UniFacePoint-FM on the TaiZhou training set. The training set contains only 3D face data without any facial attribute labels in order to prevent information leakage. An overview of the pretraining framework is shown in Fig. 1. A face is partitioned into patches, a large portion of which are randomly masked. The visible patches are embedded and fed into a Transformer-based autoencoder, which learns high-level latent facial representations from the unmasked regions and reconstructs the masked patches. To accommodate the high density and subtle geometric variations of facial scans compared with generic 3D objects, we increase both the number of patches and the patch size during partitioning. The mask ratio of 0.6 is expected to promote high generalization capability. As shown in Fig. 2, points in the masked region are well reconstructed on the TaiZhou validation and test set, similar to the results on the training set. It indicates our model is undergoing effective and stable pretraining.

**Fig. 2:**
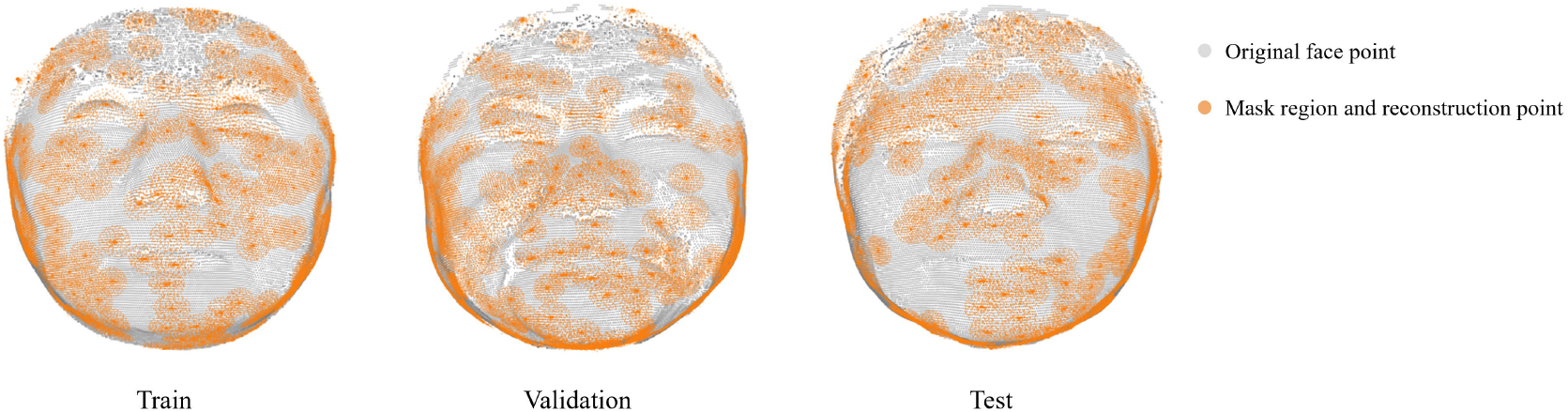
Reconstruction examples on TaiZhou train/validation/test set

### 4.2. Downstream tasks

In the pretrain stage, only 3D face data without any facial attribute labels was used, ensuring that the model learns intrinsic facial representations without task-specific biases. To evaluate the effectiveness and generalizability of the learned representations, we fine-tuned the pre-trained UniFacePoint-FM on multiple downstream tasks across several datasets.

In the TaiZhou dataset, 2,660 samples contain paired gender, age, and BMI annotations. As shown in Fig. S1(a), the gender distribution is slightly imbalanced but still adequate for gender classification, while both age and BMI exhibit wide ranges and reasonable distribution, making this dataset suitable for age and BMI prediction as well.

The ChenZhou dataset is also a self-constructed dataset, which includes 1,732 3D facial scans with corresponding gender, age, and BMI labels. However, the age distribution ranges only from 17 to 32 years and is highly concentrated around 20, making it unsuitable for age regression. Therefore, we evaluate only gender classification and BMI prediction on the ChenZhou dataset.

In addition, we incorporate the publicly available FaceScape dataset to further assess the model’s performance in gender classification and expression recognition. It is worth noting that facial scans in this dataset contain only 12,296 points—substantially fewer than the 32,251-point representations used in the other two datasets. Moreover, FaceScape exhibits a more balanced gender distribution, in contrast to the female-dominant distributions in the TaiZhou and ChenZhou datasets. Therefore, strong performance on FaceScape would further demonstrate the model’s robustness to varying point-cloud densities as well as the generalizability of its learned representations across datasets with different demographic characteristics.

#### 4.2.1. Gender Classification

To evaluate the effectiveness of UniFacePoint-FM in gender classification, we conducted fine-tuning experiments on three heterogeneous 3D facial datasets: TaiZhou, ChenZhou, and FaceScape. As illustrated by the ROC curves in Fig. 3, the model exhibits excellent discriminative performance across all test sets, with AUC values approaching 1.0. The confusion matrices further support these results, showing gender-specific classification accuracies exceeding 97% on both the TaiZhou and ChenZhou datasets. Although faces in the FaceScape dataset have fewer points and different gender distribution compared with the other two datasets, the model’s performance remains high, correctly classifying 37 males (100.00%) and 43 females (93.48%), with only three female samples misclassified. Quantitative comparisons are summarized in Table 1. UniFacePoint-FM consistently outperforms established benchmarks, including Random Forest (RF), Support Vector Machine (SVM), and ResMLP. In particular, it surpasses the RF baseline by more than 8 percentage points in accuracy on both the ChenZhou and FaceScape datasets.

**Table 1:** Performance comparison of different models on TZ-Test, CZ-Test, and FS-Test.

| Method | TZ-Test |  |  | CZ-Test |  |  | FS-Test |  |  |
| --- | --- | --- | --- | --- | --- | --- | --- | --- | --- |
|  | ACC | AUC | F1-score | ACC | AUC | F1-score | ACC | AUC | F1-score |
| RF | 91.85% | 0.9761 | 93.85% | 89.14% | 0.9670 | 92.31% | 87.95% | 0.9407 | 89.13% |
| SVM | 94.81% | 0.9891 | 96.11% | 94.29% | 0.9917 | 95.73% | 87.95% | 0.9539 | 89.13% |
| ResMLP | 95.96% | 0.9853 | 96.70% | 96.57% | **0.9950** | 97.46% | 91.57% | 0.9806 | 91.95% |
| Ours | **98.52%** | **0.9980** | **98.90%** | **98.29%** | 0.9923 | **98.70%** | **96.39%** | **0.9859** | **96.63%** |

**Fig. 3:**
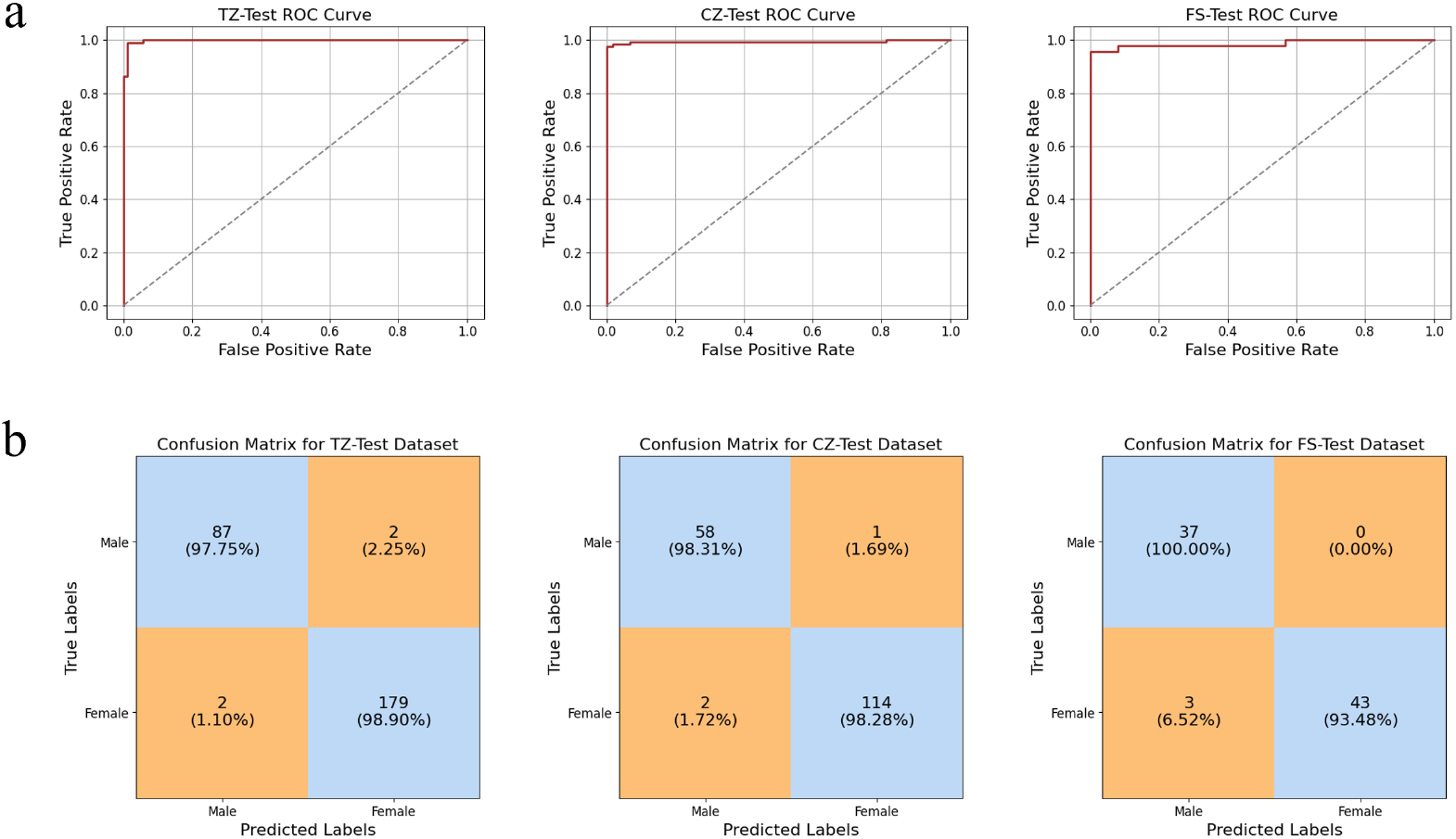
ROC curves and confusion matrices for gender classification on TZ-Test, CZ-Test and FS-Test

#### 4.2.2. Age Prediction

In this study, we evaluated our UniFacePoint-FM model for age prediction on the TaiZhou dataset. As illustrated in Fig. 4. the predicted ages exhibit a strong linear relationship with the ground truth values, with most samples closely distributed around the unity line (y =x). We further compared our method with support vector regression (SVR) and partial least squares regression (PLSR). On the TaiZhou test set, UniFacePoint-FM achieves a mean absolute error (MAE) of 3.20 and a correlation coefficient of 0.88 on the TaiZhou test set, outperforming both SVR and PLSR baselines, as summarized in Table 2.

**Table 2:** Summary of age regression performance on TZ-Test dataset.

|  | **MSE** | **MAE** | **R<sup>2</sup>** | **CORR** |
| --- | --- | --- | --- | --- |
| PLSR | 24.96 | 3.97 | 0.70 | 0.84 |
| SVR | 38.19 | 4.98 | 0.54 | 0.75 |
| **Ours** | **18.86** | **3.20** | **0.78** | **0.88** |

**Fig. 4:**
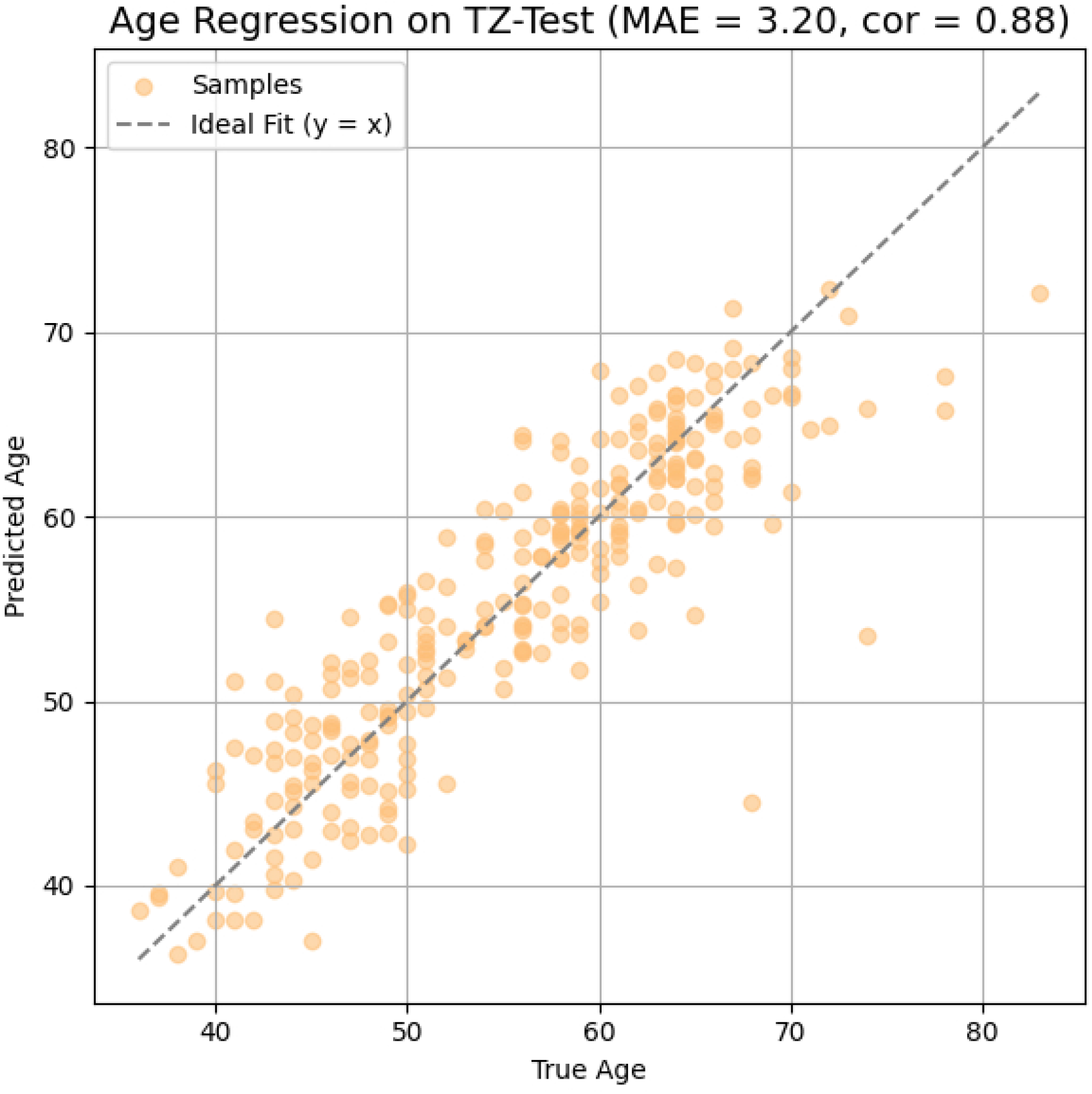
Age regression 1p6erformance on TZ-Test

#### 4.2.3. BMI Prediction

We explored UniFacePoint-FM in BMI prediction from 3D facial scans with fine-tuning strategy in both TaiZhou and ChenZhou datasets. As shown in Fig. 5, the predicted BMI values exhibit a moderate-to-strong correlation with the ground-truth measurements on both datasets. The corresponding quantitative regression results are summarized in Table 3. On the TaiZhou dataset, our model consistently outperforms the competing methods across all evaluation metrics. On the ChenZhou dataset, UniFacePoint-FM achieves the best MAE (1.28), slightly outperforming PLSR (1.31), while the remaining metrics show comparable performance among all methods. Additionally, the relatively lower MAE observed in the ChenZhou dataset compared to TaiZhou is primarily attributed to a more concentrated distribution of BMI values within the ChenZhou student population.

**Table 3:**
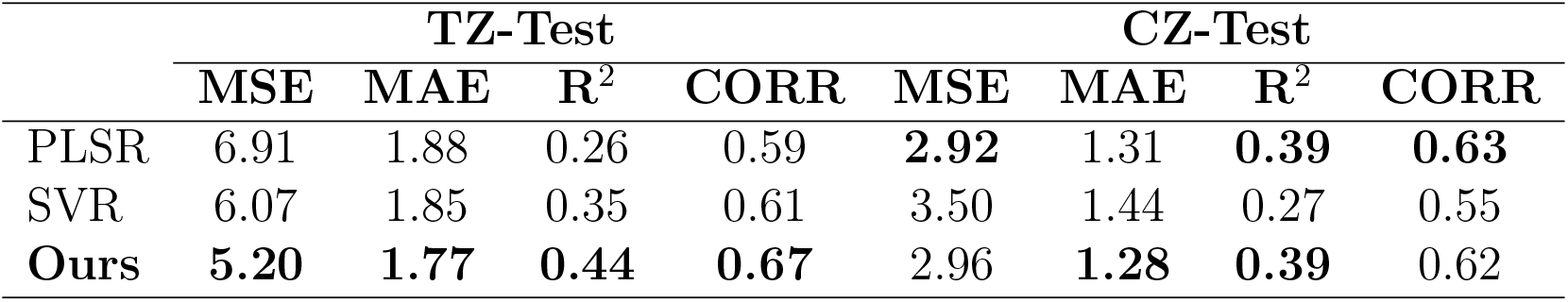
Summary of BMI regression performance on the TZ-Test and CZ-Test datasets.

|  | **TZ-Test** |  |  |  | **CZ-Test** |  |  |  |
| --- | --- | --- | --- | --- | --- | --- | --- | --- |
|  | **MSE** | **MAE** | **R<sup>2</sup>** | **CORR** | **MSE** | **MAE** | **R<sup>2</sup>** | **CORR** |
| PLSR | 6.91 | 1.88 | 0.26 | 0.59 | **2.92** | 1.31 | **0.39** | **0.63** |
| SVR | 6.07 | 1.85 | 0.35 | 0.61 | 3.50 | 1.44 | 0.27 | 0.55 |
| **Ours** | **5.20** | **1.77** | **0.44** | **0.67** | 2.96 | **1.28** | **0.39** | 0.62 |

**Fig. 5:**
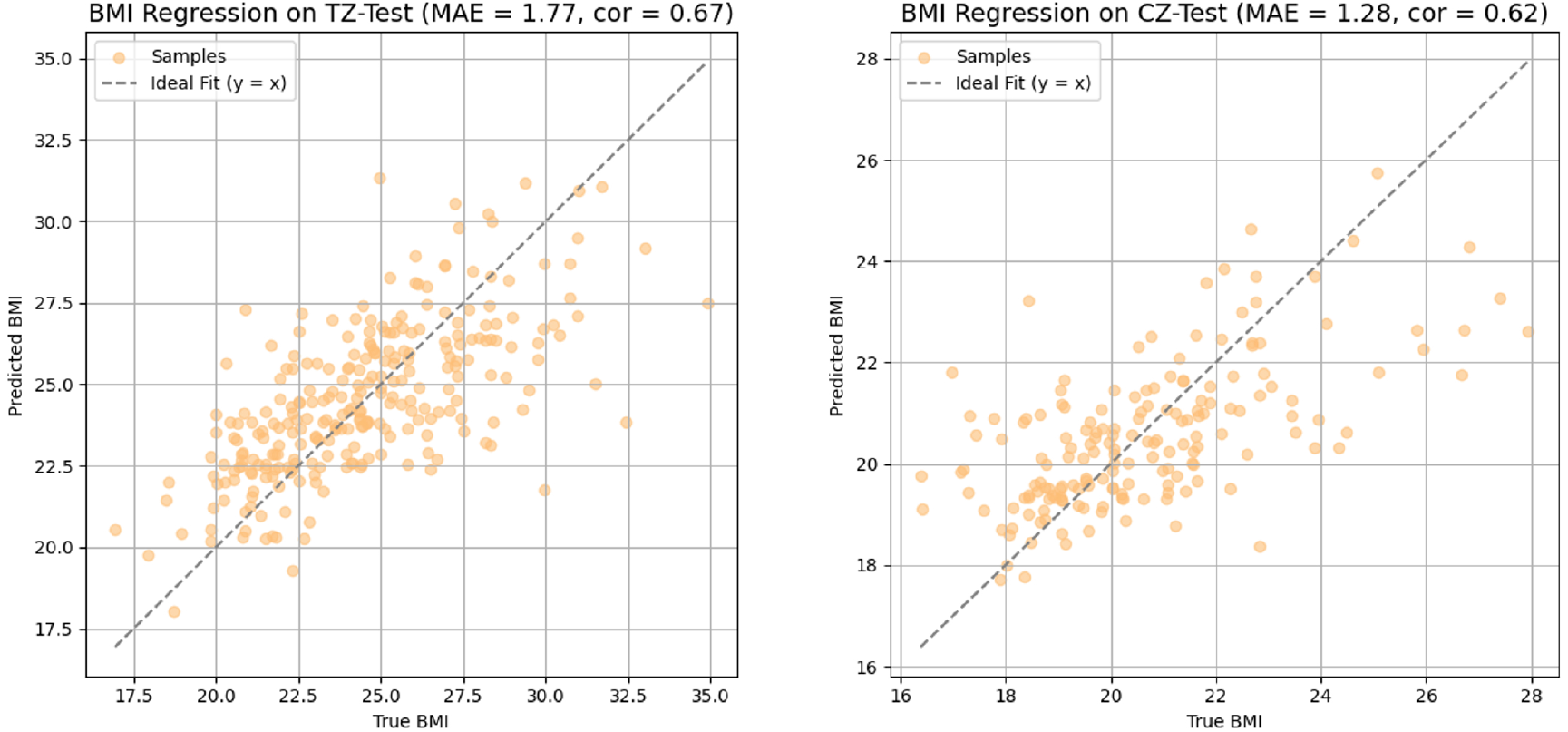
BMI regression performance on TZ-Test and CZ-Test.

#### 4.2.4. Expression Recognition

We benchmarked UniFacePoint-FM against several comparison methods on the FaceScape dataset. Fig. 6 illustrates the per-class precision and F1score radar plots for the FaceScape test set. Our method achieves consistently high performance across all expression classes, producing an almost perfectly circular contour near the upper bound of the radar chart. This near-circular contour indicates a relatively balanced performance across expression categories and stable behavior in the multi-class classification task. In contrast, SVM and RF exhibit noticeable performance degradations in specific categories, as reflected by inward-collapsed polygon shapes or prominent spikes in their radar profiles. ResMLP shows more competitive results with curves closely approaching ours. Overall, our model attains an accuracy of 99.82% and an F1-score of 99.82%, matching ResMLP and clearly outperforming the other traditional methods, as summarized in Table 4. Notably, compared with RF, UniFacePoint-FM improves the overall accuracy by 5.35 percentage points.

**Table 4:** Summary of facial expression classification performance on FS-Test.

|  | FS-Test |  |
| --- | --- | --- |
|  | ACC | F1-macro |
| SVM | 98.53% | 98.54% |
| RF | 94.47% | 94.48% |
| ResMLP | 99.82% | 99.82% |
| Ours | **99.82%** | **99.82%** |

**Fig. 6:**
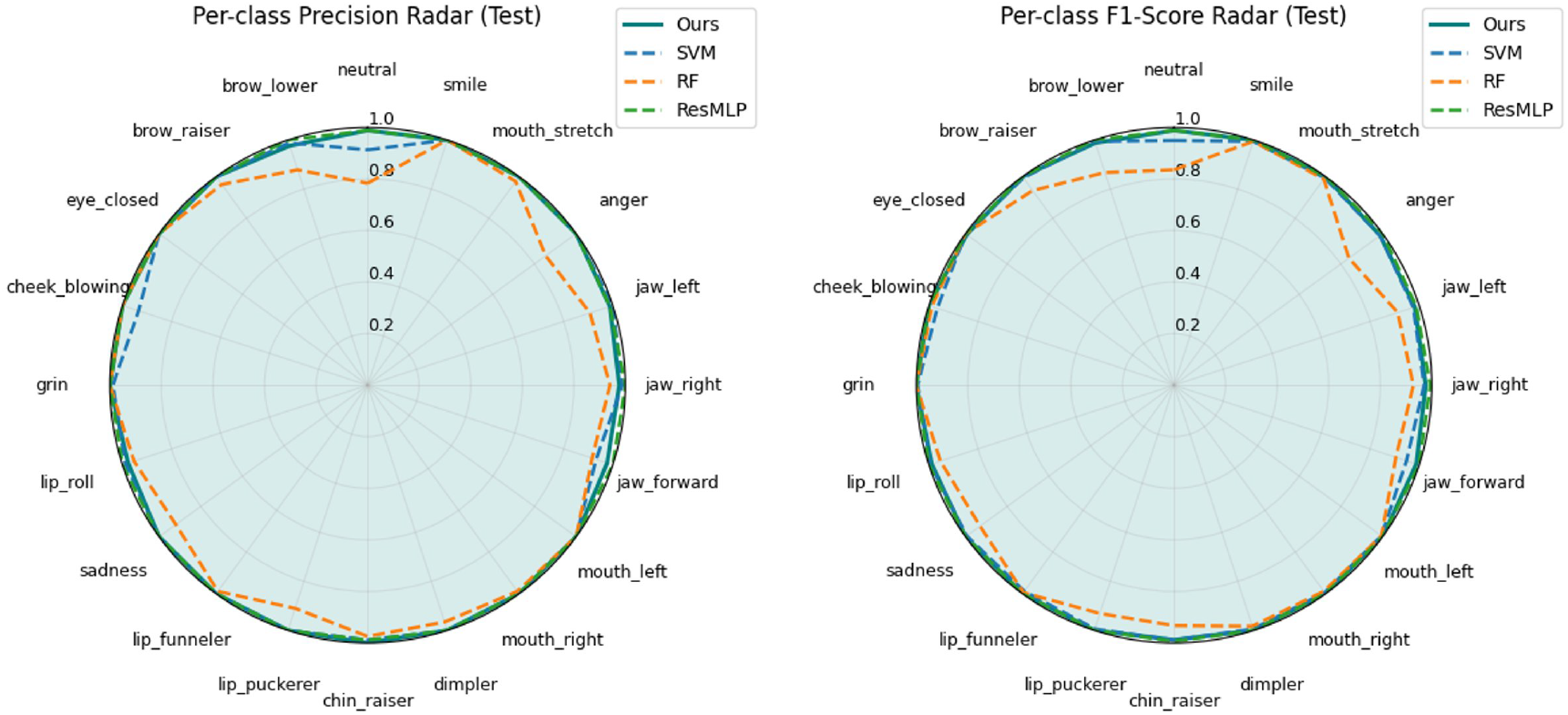
Expression classification performance on FS-Test.

## 5. Discussion

In this study, we presented UniFacePoint-FM, a framework that transitions 3D facial analysis from task-specific engineering to a more scalable selfsupervised representation learning paradigm. By utilizing a masked autoencoder architecture on large-scale unlabeled point clouds, the model captures complex geometric structures without the constraints of linear subspaces inherent in traditional 3D Morphable Models (3DMMs), or the sensitivity to environmental factors such as illumination and texture often found in 2Dbased methods.

The model demonstrated consistent performance across several distinct downstream tasks. In physiological attribute tasks, including gender classification, age regression, and BMI prediction, UniFacePoint-FM outperformed traditional linear baselines like PLSR and SVR. Specifically, in facial expression recognition, the model achieved performance comparable to state-of-the-art architectures like ResMLP, indicating its ability to represent both stable anatomical features and transient muscular deformations.

A notable observation is the model’s capacity for cross-dataset generalization. Despite being pretrained on data captured via a specific 3dMD system, the representations remained effective when applied to the FaceScape dataset, which involves different acquisition hardware and lower point densities. This suggests that the self-supervised pretraining process helps the model extract fundamental facial priors rather than overfitting to sensor-specific noise or narrow demographic characteristics. Such robustness is particularly relevant for clinical phenotyping and biometric applications where hardware standardization is often lacking.

There are limitations to the current approach that warrant further investigation. While UniFacePoint-FM is effective for feature extraction and attribute prediction, it does not yet possess the generative flexibility or semantic parameterization offered by 3DMMs for facial synthesis. In fact, as a pure neural network-based framework, a generative expansion of UniFacePoint-FM could potentially offer more coherent and robust facial rendering than traditional 3DMMs by bypassing their linear constraints and capturing more complex geometric manifolds. Additionally, while this work utilized task-specific fine-tuning, developing a unified multi-task architecture could further improve computational efficiency and exploit the biological correlations between different facial traits. In summary, UniFacePoint-FM provides a generalizable and scalable foundation for 3D facial representation, offering a robust tool for future research in digital health and spatial intelligence.

## Supplementary Material

**Fig. S1:**
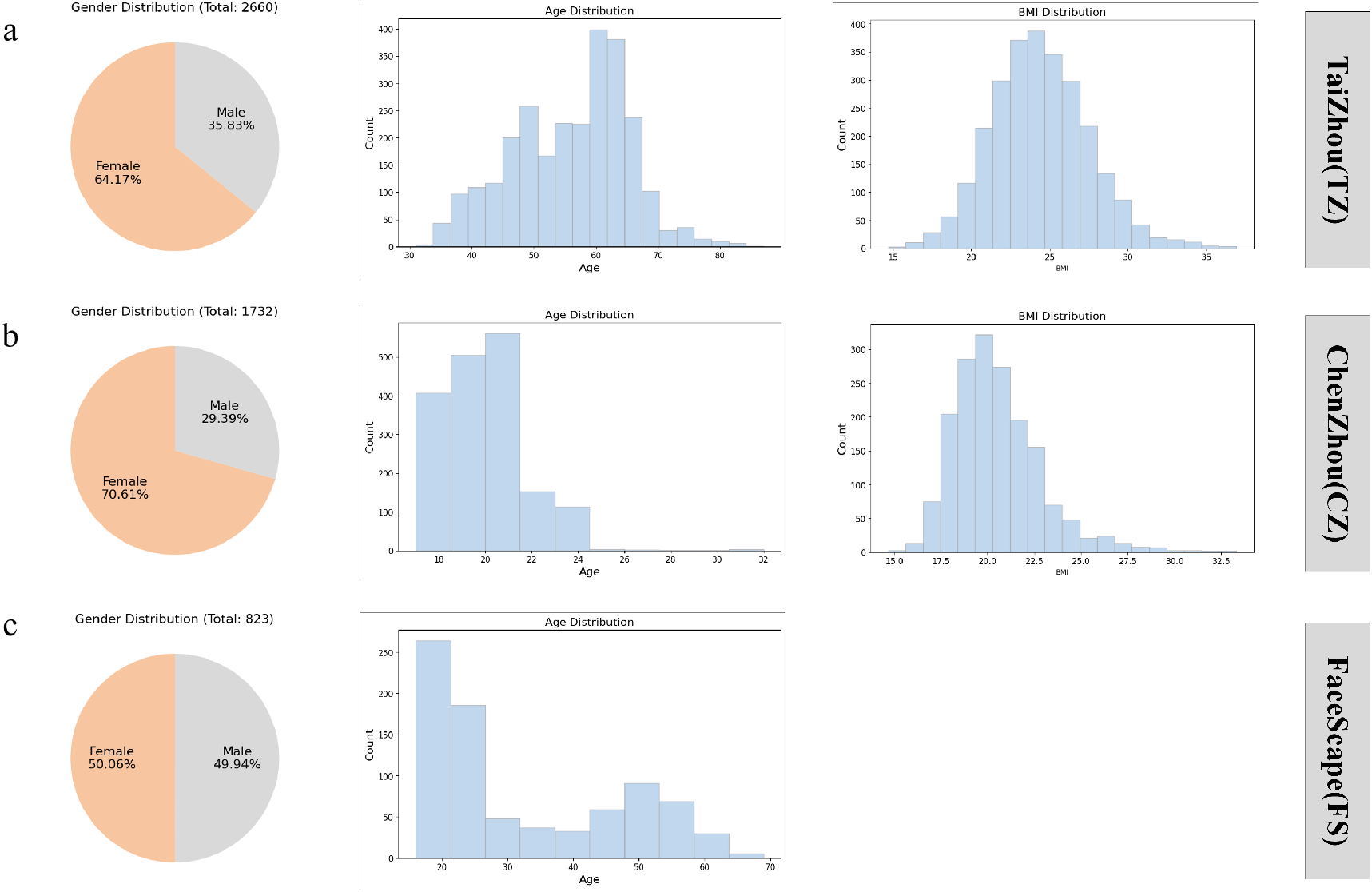
Demographic distributions of participants across three databases.

